# A Multiscale Computational Analysis of Myometrial Excitation during Late Pregnancy

**DOI:** 10.64898/2026.06.22.733909

**Authors:** Parker Roman Mixon, Vijay Vedula

## Abstract

The control of uterine activity during pregnancy is a complex process that involves regulating myometrial excitability across multiple scales. While numerous studies have investigated various regulatory mechanisms and established the contributions of ion channels and gap junctions, how these mechanisms interact to produce observed changes in uterine activity remains poorly understood. Pivotal to these efforts are computational models that effectively capture gestational changes in excitability across scales. In this study, we propose a multiscale computational modeling framework that can reproduce measured activity at the cellular and tissue scales at a given gestational stage. At the cellular level, we identify key ion currents underlying the observed electrophysiological properties based on a literature review of their regulation and a sensitivity analysis of the Tong 2011 uterine smooth muscle cell activation model. The conductances of these ion currents are then fit to reproduce characteristic resting membrane potentials and burst properties using Bayesian optimization. To extend to the tissue level, we employ an anisotropic monodomain model, parameterized by the resistivity of late pregnancy uterine muscle, to investigate electrical propagation in a two-dimensional section of uterine tissue. We then apply the multiscale model to study myometrial activation in late pregnancy and elucidate the contributions of ion channel and gap junction regulation in transitioning the uterus from a quiescent state to labor. Our resulting model successfully reproduces measured electrophysiological properties at the cellular level and characteristic single-spike and burst-propagation patterns at the tissue level across the three late-pregnant time points analyzed (days 16/17, 18/19, and 20/21) in a murine model. Furthermore, our results suggest that the regulation of the conductances of the voltage-dependent potassium current (I_K1_), L-type calcium current (I_CaL_), and sodium current (I_Na_) is most important in determining preterm uterine excitability. The framework established here will promote the development of more gestationally relevant models to better understand labor progression and the factors involved in dysfunctional labor.

**Author Summary:** Pregnancy is marked by drastic changes in the electrical and contractile activity of the uterus. As improper regulation of uterine activity is associated with preterm and dysfunctional labor, it is crucial to understand the physiological mechanisms underlying these changes. Currently, the roles of ion channels in determining cellular dynamics, and gap junctions in cell-to-cell coupling, as well as tissue properties, have been well established. However, how their regulation interacts to produce observed changes in cellular and tissue excitability and, in turn, organ-level activity is far less understood. While existing computational models of uterine electrophysiology have provided a greater insight into these processes, these are formulated for a single time point and cannot interrogate their effects over gestation. In response to this need, we develop a framework to generate a computational model of uterine excitation at a given gestational stage. We apply this to investigate the role of ion channels and gap junctions in transitioning the uterus to labor during late pregnancy. We identify three major ion channels and demonstrate agreement with observed action potential and tissue propagation properties at the analyzed time points. We further highlight how our framework can be applied to investigate other stages, including labor and postpartum.

## 1. Introduction

Careful control of myometrial activity is necessary for the safety of the developing fetus and successful delivery at term. As the term approaches, the uterus transitions from a quiescent state, prevalent for most of gestation, to a highly contractile state, capable of producing the forceful, coordinated contractions required for delivery of the fetus [1]. This process occurs throughout gestation via the regulation of ion channels and other contraction-associated proteins, but is most significant in late pregnancy and labor, where it is known as *myometrial activation* [2,3]. Early activation of the myometrium is one of the major contributors to preterm birth, which affects nearly 11% of deliveries worldwide and is the most common cause of neonatal deaths [4]. On the contrary, incomplete or delayed myometrial activation may result in abnormal labor progression termed *dystocia* [4]. Dystocia remains the primary indication for cesarean delivery and is associated with an increased risk of maternal mortality and morbidity [5]. Our objective in this study is therefore to unravel the mechanisms regulating myometrial quiescence and activation during late pregnancy, using a novel multiscale computational model that incorporates gestational changes in uterine smooth muscle cell (USMC) and tissue excitability. Understanding these pathways is critical for assessing the risks of labor complications, such as preterm labor, as well as for identifying potential therapeutic targets.

As the term approaches, changes in uterine excitability and electrophysiological (EP) properties can be observed across multiple scales. At the cellular level, action potentials (APs) transition from single spikes to long-lasting bursts, increasing cellular force generation, while the resting membrane potential (RMP) depolarizes from -60 mV in the non-pregnant uterus to -45 mV at term [6]. These transformations occur primarily due to changes in ion channel expression and activity, although the exact contributions of individual channels have proven difficult to identify, especially over the course of gestation [6]. At the tissue level, the upregulation of gap junctions provides low-resistance pathways between USMCs, allowing APs to propagate farther and at faster velocities [7]. However, studies have shown that an increase in gap junctions alone does not fully explain the transition to the highly synchronized activity observed during labor.

This is further hindered by the fact that the human uterus lacks a specific pacemaker, and, as such, the origin of excitation is unknown [7,8]. Instead of a designated anatomical pacemaker, as seen in the heart, experiments have shown a possible role of mechanotransduction in producing organ-level coordination. In this theory, the increase in intrauterine pressure from contractions deforms the uterine wall, triggering stretch-activated contractions in other areas of the wall [9]. Another proposal is that recently discovered interstitial cells, called *telocytes*, may form a pacemaking network analogous to that of the interstitial cells of Cajal in the stomach [2]. In the uterus, however, these cells have instead been shown to hyperpolarize and may therefore play a role in maintaining quiescence [2].

While studies have generated numerous hypotheses regarding potential mechanisms that regulate labor and uterine contractility, such as those mentioned above, it has been challenging to verify their roles in human pregnancy using experimental methods. Moreover, ethical considerations restrict the use of many imaging and experimental methods on humans [10]. The applicability of data from animal models is limited by key physiological differences, including uterine structure, offspring number, and the method of progesterone withdrawal [10]. In response to these limitations, electrohysterography has emerged as a promising research tool for noninvasive imaging of uterine activity in humans; however, further work on signal-processing, standardization, and model validation is required before widespread clinical adoption [11,12].

Computational modeling, on the other hand, being noninvasive and predictive, forms the cornerstone of pregnancy research, overcoming the challenges associated with *in vivo* and *in vitro* approaches. Single-cell models of USMC excitation based on biophysically-detailed ion current formulations have provided valuable insights into the physiology of AP generation and excitation-contraction coupling, thereby supporting the roles of ion channel and gap junction regulation in myometrial activation [13–17]. Conversely, tissue- and organ-level models have investigated the origins of uterine activation and synchronization during labor, although they often employ simplified or phenomenological models of cellular activation that do not account for specific ion channel contributions or their modulation across gestation [18–22]. A recent computational study by Yang et al. has overcome many of these limitations by coupling the Tong 2014 cellular activation model [15] with an anisotropic diffusion model at the organ level [23].

However, as with many other models, their investigation primarily focused on the term uterus, although the effects of uterine shape and tissue conductivity on activation patterns were also analyzed. Despite these computational advances, there remains an unmet need to synthesize the various debated mechanisms at the cellular, tissue, and organ levels into a cohesive, multiscale model of myometrial activation and labor progression.

To this end, we identify a few limitations of existing models. First, they are formulated for a single time point and cannot account for differing activities across gestational stages nor for the progression of excitability during labor. This limits their application to the key use case of predictive modeling of preterm labor and other labor risks, including dystocia. Second, current models lack sufficient physiological detail across the cellular-, tissue-, and organ-scales to effectively interrogate the role of multiscale interactions and modulation in determining uterine activity. Our study, therefore, represents a first step toward addressing these limitations at the cellular and tissue levels, with a primary application to the study of myometrial activation in the preterm uterus. Through a multiscale computational analysis of myometrial activation, we aim to both validate our approach’s ability to reproduce experimental AP and tissue propagation properties and to elucidate the role of ion currents and gap junctions in transitioning the pregnant uterus from late gestation to labor.

## 2. Results

### 2.1. Sensitivity analysis

The first-order Sobol’ indices show the direct effect of a single current’s conductance value on each measured EP property (Figure 1A). From these first-order indices, we see that only the non-selective cation current (I_NSCC_), background current (I_b_), I_CaL_, and I_K1_ currents had any substantial impact alone on the model’s EP properties, with the strongest effects being on the RMP, followed by the height and the rate of rise of the initial spike (Figure 1A). The spike density and the burst duration showed less sensitivity (below an index of 0.05) to changes in a single current’s conductance. On the other hand, the total-order Sobol’ indices, given in Figure 1B, account for both the first-order effects of a current and its interactions with all other currents in affecting the measured property. The total-order indices for all the burst properties, including height, rate of rise, burst duration, and spike density, were much greater than the first-order indices, suggesting that bursts are the result of complex interactions between many currents. The RMP, in contrast, was mainly sensitive to the I_NSCC_, I_K1,_ and I_b_ currents, showing little change between first- and total-order indices (Figure 1B). The burst properties were most sensitive to the I_NSCC_, I_K1_, I_b_, I_CaL_, I_CaT_, and I_Na_ currents, although they remained only moderately sensitive to many of the other currents.

**Figure 1.**
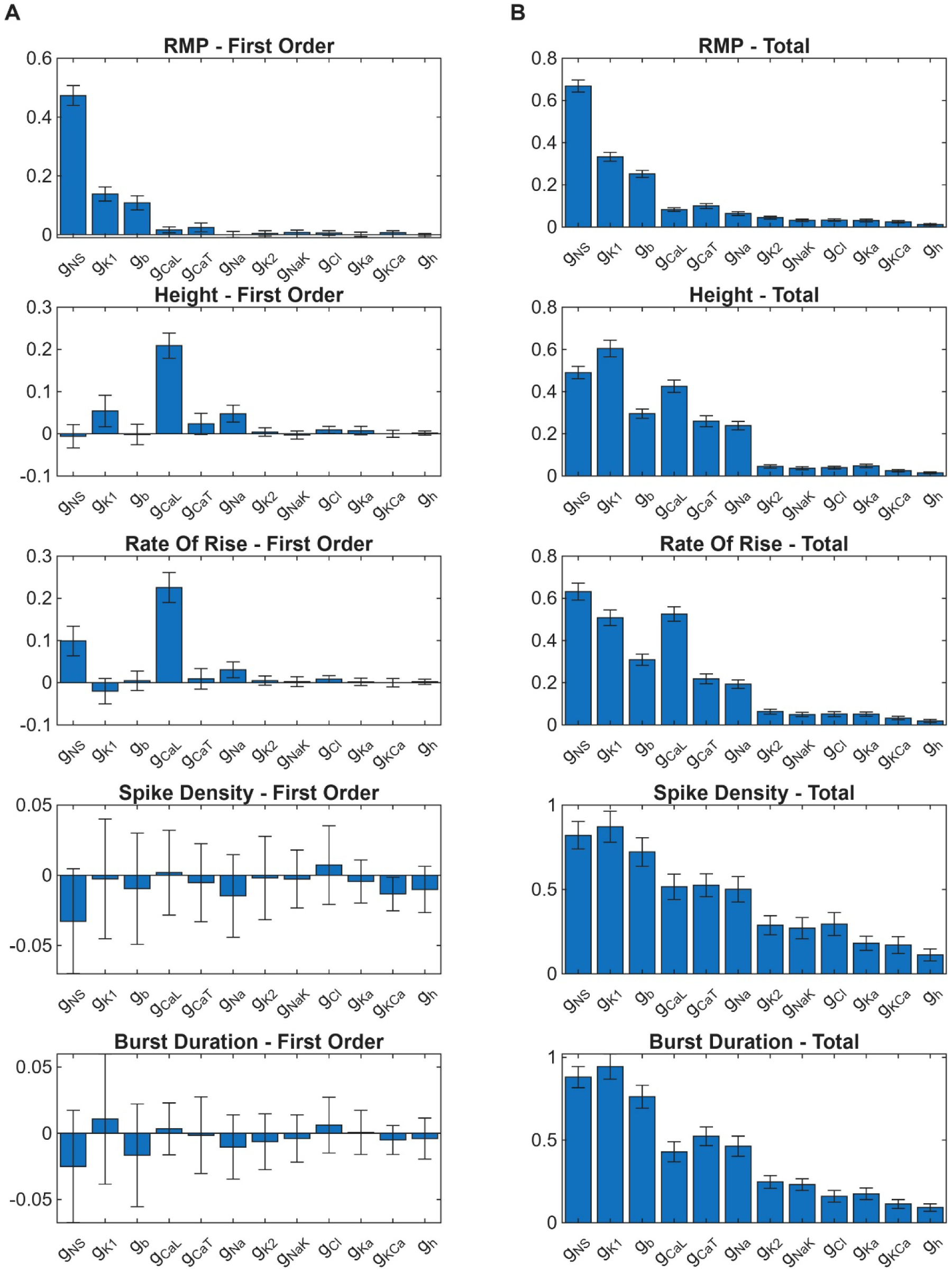
Global variance-based sensitivity analysis of the Tong model. First- and total-order Sobol’ indices of the ion currents of the Tong 2011 model evaluated using 4096 samples. All ion currents are sorted in descending order of the sum of their total-order indices. Error bars represent the 95% confidence interval computed by bootstrapping. Only 12 of the 14 ion currents in the Tong 2011 model are represented here. The hyperpolarization current (I_h_) was assumed to be constant due to the absence of literature on its gestational regulation, while the Na^+^-Ca^2+^ exchanger current (I_NaCa_) was excluded as it lacks a maximal conductance parameter and instead depends on the corresponding Na^+^-Ca^2+^ flux (J_NaCa_). (A) First-order Sobol’ indices representing the effect of the particular ion current alone on the measured electrophysiological (EP) properties. (B) Total-order Sobol’ indices representing the combined effect of the specific ion current with all other currents on the measured properties.

### 2.2. Best-fit conductance values and cellular electrophysiological properties

Using Bayesian optimization, the maximal conductances of the I_K1_, I_CaL_, and I_Na_ currents were fit to reproduce characteristic RMP, burst duration, and spike density values at three time points in late pregnancy. The resulting EP properties agreed reasonably well with the literature values (Figure 2A), with total relative errors of 1.29%, 0.99%, and 1.63% for days 16/17, 18/19, and 20/21, respectively. The corresponding maximal conductance and stimulus values generally showed an increase in the conductances of the excitatory currents I_Na_ and I_CaL_ and a decrease in the conductance of the inhibitory current I_K1_ with gestational age (Figure 2B). However, day 18/19 appeared to be an outlier with a substantial drop in both I_CaL_ and I_K1_ conductances. The stimulus amplitude also decreased with gestational age, likely in response to the increased cellular excitability. This increase in cellular excitability with gestational age is also readily apparent from the resulting burst APs (Figure 2C), where bursts at the preterm time points, days 16/17 and 18/19, gradually increased in amplitude following the initial spike. In contrast, the term time point (day 20/21) maintained consistent amplitudes throughout the burst sequence.

**Figure 2.**
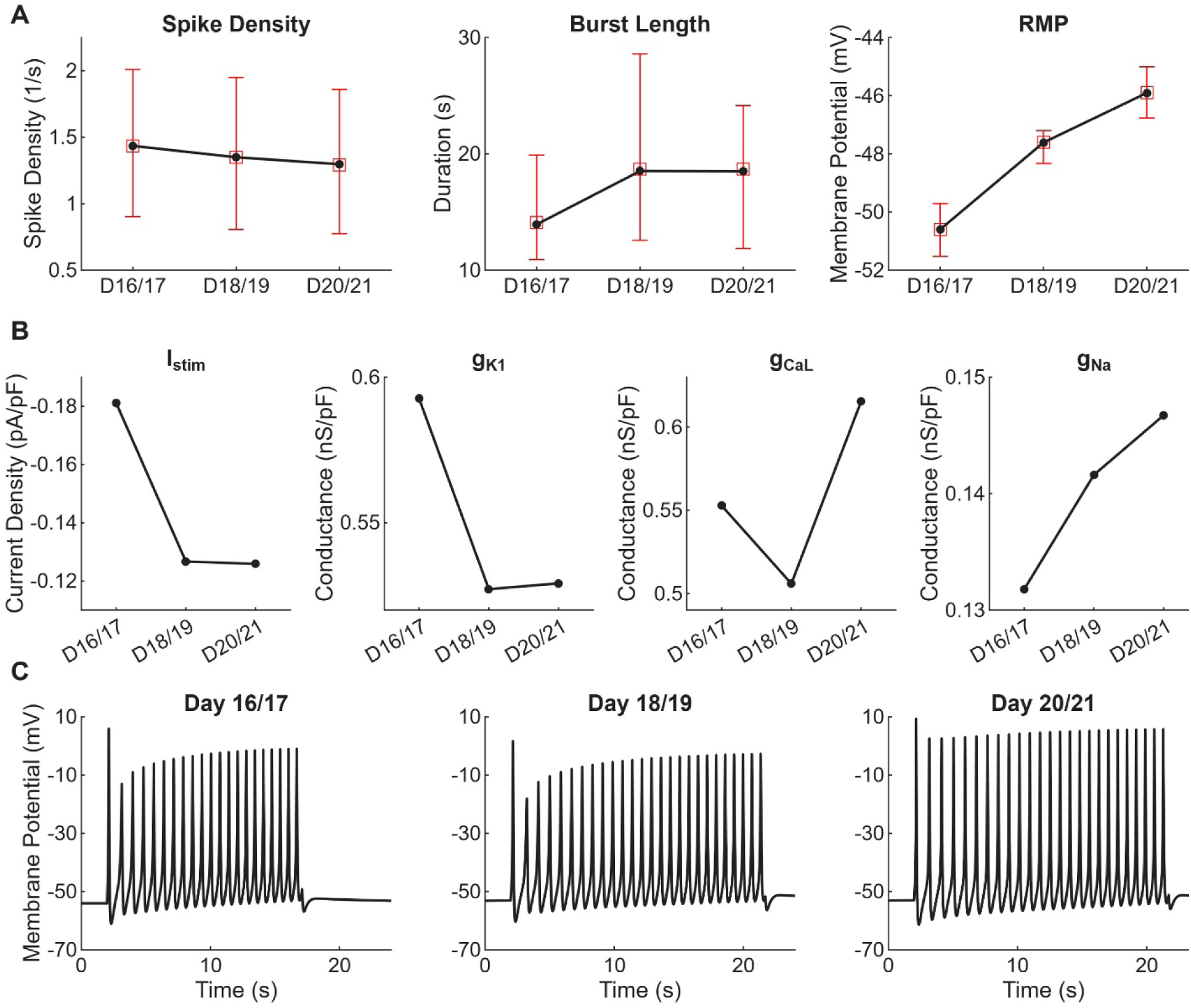
Best-fit parameters and trends for three time points in late pregnancy obtained using Bayesian optimization. (A) Properties of best-fit burst compared to corresponding literature values. Best-fit values are shown as black circles, while the red squares and error bars are from the respective literature data. For the spike density and the burst length, the red square represents the median, and the error bars represent the interquartile range from Reinl et al. [24]. For the resting membrane potential (RMP), the data represent the mean ±1 standard deviation from Bengtsson et al. [25]. (B) Trend of best-fit conductance and stimulus values. (C). Best-fit gestational bursting-type APs using default extracellular concentrations.

### 2.3. Single-spike and burst propagation

To investigate the properties of AP propagation in uterine tissue, we simulated single-spike and bursting APs on a two-dimensional plane at each gestational time point, using the simulation protocol provided in Fig. A2 of the Supplemental Material. In all cases, the spike propagated throughout the domain in a highly anisotropic manner, with faster conduction along the longitudinal axis (vertical, y-axis). The conduction velocities measured from the activation map for a single-spike AP (Figure 3) are given in Table 1 for both the longitudinal and circumferential directions. The recorded velocities are in reasonable agreement with literature values [7]. Although the diffusion coefficient was identical across gestational time points, the conduction velocities were substantially different, likely reflecting variations in cellular excitability. In general, conduction velocity increased with gestational age; however, day 18/19 again appears as an outlier, exhibiting the slowest velocity.

**Figure 3.**
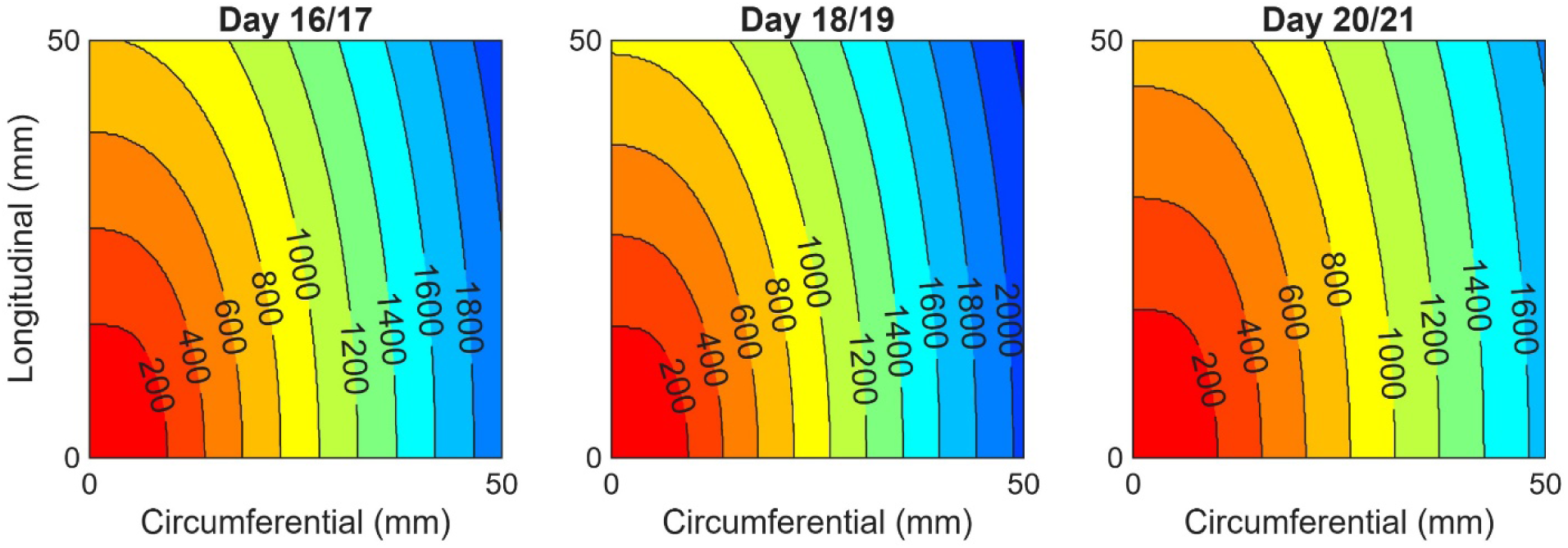
Single-spike activation maps across three time points in late pregnancy. Single spikes were evoked by a stimulus of -5.0 pA/pF for 50 ms to a 10 mm square area in the bottom-left corner of the domain. Isochrones are plotted at a 200 ms interval. Grid spacing (Δx = 0.25 mm) and time step size (Δt = 0.5 ms) were determined from a convergence analysis (Sec. S1 of Supplemental Material).

**Table 1.** Single spike conduction velocities obtained from the activation maps shown in Figure 3.

| Conduction Velocity (cm/s) |  |  |
| --- | --- | --- |
| Gestational Day | Longitudinal | Circumferential |
| 16/17 | 5.78 | 2.34 |
| 18/19 | 5.45 | 2.20 |
| 20/21 | 6.69 | 2.71 |

**Figure 4.**
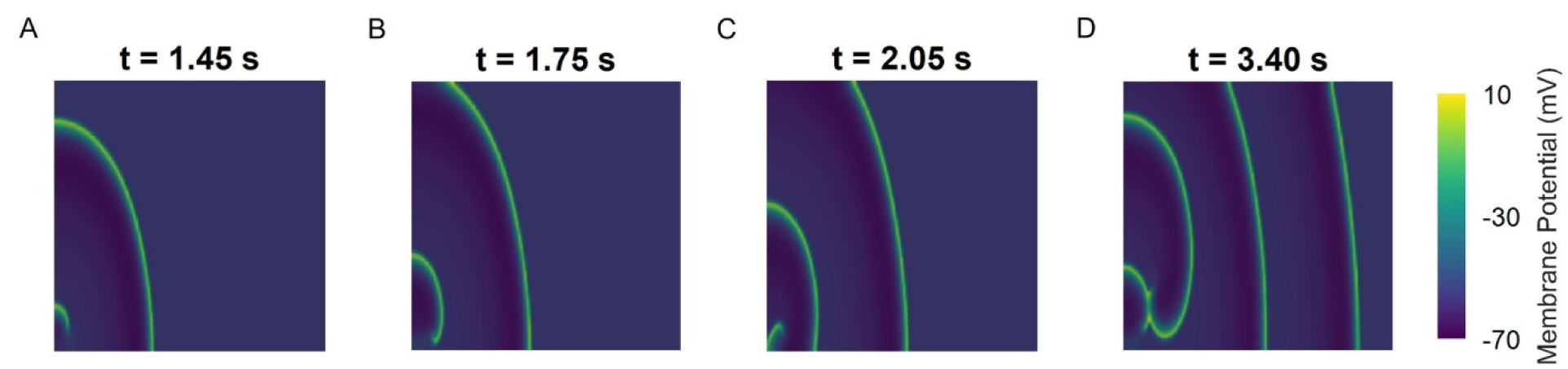
Example of spiral waves observed for burst propagation in the day 20/21 tissue model. Propagation of a burst evoked by a stimulus of -0.15 pA/pF for 18.7s to a 6 mm square area in the bottom left corner of a 10 x 10 cm two-dimensional domain. (A-C) The development of a spiral wave after the initial spike fails to propagate circumferentially through the length of the domain. (D) The collision of the initial spiral wave with a newly developed second wave. Grid spacing (Δx = 0.25 mm) and time step size (Δt = 0.5 ms) were determined from a convergence analysis (Sec. S1 of Supplemental Material).

While the initial spikes of the resulting burst frequently propagated throughout the tissue, subsequent spikes occasionally failed to propagate, especially with low-amplitude stimuli or smaller stimulation areas (**Error! Reference source not found.**A). Progressive recruitment, as described by Lammers for day 17 rat myometrium, was not observed [26]. When spikes failed to propagate circumferentially (x-axis) but continued longitudinally (y-axis), reentrant waves were occasionally seen. Since the spikes traveled through the entire tissue, these instances of re-entry often developed into long-lasting spiral waves. A representative reentrant wave is shown for day 20/21 in **Error! Reference source not found.**.

## 3. Discussion

Despite numerous proposed mechanisms, the control of uterine activity during pregnancy remains poorly understood, although it is known to arise from the complex interaction and regulation of processes across multiple scales [2]. Computational models have proven useful for investigating these mechanisms. Cellular models have provided insight into the contributions of ion channels to AP properties and force generation, while tissue- and organ-level models have studied the origin and control of uterine activity [13–23]. However, a key limitation of these models is that they are formulated for a single gestational time point and often lack sufficient physiological detail across scales, thereby limiting their ability to predict normal and preterm labor progression.

To address these limitations, we have developed a framework to generate a multiscale model of the uterus capable of reproducing EP dynamics at various gestational time points by incorporating corresponding adaptations in ion channels and gap junctions at the cellular and tissue levels. We then applied these methods to investigate myometrial activation in the late-pregnant uterus. At the single-cell level, we developed a methodology to fit USMC activation models to different gestational time points. We demonstrated it using the widely used Tong 2011 model to reproduce characteristic changes in EP properties observed in late pregnancy. At the tissue-level, we employed a monodomain model with anisotropic diffusion to investigate the effects of ion channel and gap junction regulation on AP propagation. Overall, our model reproduced SMC activation properties to within 2% relative error at all time points and conduction velocities within the ranges reported in literature for both the longitudinal and circumferential directions.

Beginning with a variance-based global sensitivity analysis at the cellular level, we examined the sensitivity of various burst properties and of the RMP to changes in ion channel conductance. We found that only the RMP and the height and rate of rise of the initial spike were sensitive to variations in a single current’s conductance, with the I_NSCC_ and I_CaL_ currents having the most substantial effects. In contrast, the duration and spike density of a burst-type AP were only affected when multiple conductances were varied, exhibiting no first-order sensitivities.

This result is consistent with the one-at-a-time sensitivity analysis performed by Tong et al. in their follow-up study, in which they were unable to extend the burst duration with a single change in channel conductance or gating parameters [15]. Burst phenomena in uterine tissue, therefore, appear to be a complex interaction between multiple currents with equally complex regulation.

Based on our sensitivity analysis and a literature review of the gestational regulation of uterine ion currents, we chose to fit the maximal conductances of the I_CaL_, I_Na_, and I_K1_ currents along with the stimulus amplitude. Using Bayesian optimization, these parameters were successfully fit to reproduce changes in RMP, burst duration, and spike density at three late-gestational time points in a murine-based model. In general, the resulting best-fit parameters showed an increase in the excitatory I_CaL_ and I_Na_ currents and a decrease in the stimulus amplitude and the inhibitory I_K1_ current. These results support the idea that the increase in uterine activity arises from a downregulation of inhibitory currents and an upregulation of excitatory ones [6].

Many of the limitations of our single-cell model arise from the limited knowledge of the gestational regulation of uterine ion channels. Existing literature is sparse and often contradictory, and no consistent timelines or values could be drawn. This precluded the direct use of recorded maximal conductances to adapt our model to each gestational time point, and therefore, a fitting procedure was required. Furthermore, data on gestational changes in ion currents in humans are limited; therefore, the model presented here is largely based on murine data. While a combination of murine, guinea pig, and human data was used to identify the most gestationally regulated ion currents, the data used for the fitting procedure are from murine models only. The use of such hybrid models that integrate data from various animal models, including the Tong 2011 model, is necessary until more human-specific data becomes available. In that case, the methods presented here can be applied directly.

The decision to model gestational ion current regulation as changes in their maximal conductances also arises from inadequate data availability, and likely does not reflect the true physiological mechanisms. Ion current regulation is highly complex, encompassing processes ranging from mRNA transcription to post-transcriptional modifications, membrane trafficking, and interactions among channel subunits. Furthermore, experimentally measured ion currents may arise from contributions of multiple ion channels; these are known as macroscopic currents. Such is the case for the uterine I_K1_ current, whose channel identities are unknown, but have been posited to include hERG, KCNQ, or Kv2.1 channels [15,16]. Indeed, in their 2014 follow-up paper, Tong et al. found that the burst duration could be prolonged by using a higher conductance I_K1_ current with slower dynamics. Based on these results, they partially replaced the I_K1_ current with hERG and KCNQ channels and successfully generated long-duration, labor-like bursts [15]. While we did not include such currents in our model, their regulation likely plays a significant role in myometrial activation and warrants further investigation. However, this would be computationally difficult with the present methods, likely requiring fitting an additional three to four currents. Future work should aim to accelerate the fitting process and investigate more complex potassium channel regulation.

To this end, it may be possible to adapt the methods of Atia et al. [16], who generated a novel activation model by fitting the conductances of an array of potential USMC ion channels, identified by transcriptomic analysis, to produce a prescribed voltage waveform [16]. While this method offers more robust fitting and a direct relationship between cellular excitability and individual ion channels rather than macroscopic ion currents, it was not used here because we could not find adequately detailed, published AP and calcium tracings at multiple gestational time points. Furthermore, the model they produced using these methods is highly complex, comprising 21 ion channels versus the 14 in the current model, and additional reduction would likely be necessary to permit extension to full organ simulation. We therefore see how these methods could be used to supplement our own, either to perform a model reduction or to refine the selection of best-fit conductances, both being informed by a combination of sensitivity analysis, gestational regulation, and functional redundancy.

We extended our single-cell activation model to the tissue level using a monodomain model of excitation-propagation, which has been widely adopted to simulate conduction in the uterus and other organs such as the heart [23,27–29]. Although we used an identical diffusion coefficient across all gestational time points, corresponding to the single late pregnant resistivity value reported by Sims et al. [30], we observed a substantial increase in conduction velocity between the preterm and term time points along both the longitudinal and circumferential fiber directions. We therefore find that this trend is associated with an increase in cellular excitability with gestational age rather than upregulation of gap junctions. The day 18/19 time point, however, appeared as an outlier, exhibiting the slowest conduction velocity, possibly corresponding to it having the lowest I_CaL_ conductance fit. This decrease in cellular excitability at day 18/19 may be a compensatory response to an upregulation of other excitatory factors (e.g., contraction-associated proteins) or to stretch-induced contractions from the growing fetus. Indeed, both the withdrawal of progesterone and the decrease in uterine growth occur around this time [31,32].

Burst propagation exhibited similar anisotropic conduction for the initial spike as that of the single spikes. Subsequent spikes, however, occasionally failed to propagate circumferentially. In these cases, reentrant phenomena were observed at all time points, consistent with Lammers et al., who reported such circulating waves in rat myometrium at days 17, 19, and 21 [33]. Our results, along with those of Xu et al. for a discrete tissue model, show that re-entry is likely inherent to burst propagation in uterine tissue and may contribute to the chaotic excitation patterns seen in labor [34]. As we were able to recreate these patterns within a continuum framework, rather than the discrete one used by Xu et al., we can readily extend our framework to a full organ model to investigate the effects of nonlinear propagation on uterine contractility and function, which will be the subject of future work.

A key limitation of our tissue model is that the APs traversed the entire domain in a highly linear fashion. We believe this limitation arises from the modeling assumptions made here, as well as in other existing models. Beginning with our use of a diffusion-based excitation-propagation model, the literature indicates that the propagation of single spikes does not occur as linear diffusion; instead, it exhibits chaotic patterns with varying conduction velocities and directions [7,8]. In fact, it is thought that AP diffusion accounts for only short-distance propagation in the uterus, while intrauterine pressure generation leading to mechanotransduction is responsible for long-distance propagation [9]. Secondly, uterine muscle tissue is likely not homogeneously excitable. There is evidence that many ion currents, such as I_Cl_, I_Na_, I_K1_, and I_K2_, are differentially expressed among individual USMCs [35–37]. The inclusion of electrically-passive cells and varying levels of heterogeneity in intercellular coupling, RMP, and cell capacitance has also been found to modulate excitability and synchrony in computational uterine tissue models [34,38]. The addition of tissue heterogeneity in a future model may therefore help to limit the propagation distance of single spikes and more effectively replicate the chaotic activation patterns observed *in vivo*. Thirdly, our assumption that fiber directions lie uniformly along two orthogonal axes (i.e., longitudinal and circumferential) is in direct contrast to the complex, interwoven fiber structure observed in a real uterus [39]. While Zhang et al. sought to address this by using random fiber orientations in their model, APs still propagated over long distances, and a manual range limit had to be set [20]. No established method exists to incorporate realistic fiber directions into uterine models, which is further hindered by the fact that uterine fiber architecture changes drastically with gestational stage [10].

Finally, as with many other models, we primarily focus on the longitudinal layer, which is the largest portion of the myometrium and is accordingly the most studied. The circumferential layer, though smaller, appears to exhibit distinct ion channel regulation and EP properties throughout pregnancy [25,40,41]. It is likely that the interaction between these muscle layers with differing excitabilities will affect the activation dynamics of the gravid uterus. In support of this idea, a recent study by Lutton et al. showed that electrical activity in the rat myometrium often originates from bridge fibers that connect the two muscle layers [42].

In conclusion, we have established a workflow to generate multiscale models of USMC excitation and propagation at various gestational time points by identifying and incorporating corresponding adaptations at both the cell and tissue levels. We applied these methods to develop a model of myometrial activation, and demonstrated its ability to reproduce characteristic EP properties, including RMP, burst duration and density, and conduction velocities at three time points in late pregnancy. In doing so, we found that modulation of the I_K1_, I_CaL_, and I_Na_ currents played a significant role in determining preterm uterine activity, highlighting them as potential targets for treating preterm labor. Gap junctions were found to play a lesser role. We have also identified possible modifications to improve fitting performance and physiological relevance, as well as the limitations of current modeling approaches. While we have focused specifically on myometrial activation in late pregnancy, the proposed methods are broadly applicable to other gestational stages, including labor and postpartum.

## 4. Methods

### 4.1. Single-cell model parameter reduction

As a baseline model for describing the excitation of individual USMCs, we employed the biophysically detailed Tong 2011 model, which has been shown to reproduce a variety of AP morphologies and hormonal effects [14]. The model describes the change in the transmembrane potential (*V_m_*) due to ion exchange through a total of 14 ion currents (*I_ion_*) using a system of differential-algebraic equations according to the Hodgkin-Huxley-type formulation:

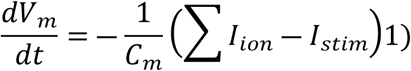

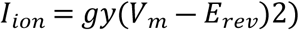

where *C_m_* is the specific membrane capacitance, *I_stim_* is the stimulus current applied to the cell, *I_ion_* is the current across a particular ion channel, *g* is the corresponding maximal conductance, *y* is the gating variable, and *E_rev_* is the reversal potential.

However, this model is also highly complex, comprising about 20 gating variables and two state variables, including *V_m_* and intracellular Ca^2+^ concentration [*Ca*^2+^]*_i_*, yielding 105 equations in total [14]. As such, we first sought to identify a subset of currents and parameters sufficient to reproduce the changes in EP properties observed in late pregnancy. We based this parameter reduction on two factors: (i) the magnitude of the gestational regulation of the currents and their properties during late pregnancy, and (ii) their influence on the EP properties of the Tong 2011 model, specifically the RMP and burst properties, such as the burst duration and spike density.

#### 4.1.1. Literature review of ion current gestational regulation

To identify the most gestationally regulated ion currents in the Tong 2011 model, we conducted a literature review for published data on changes in ion channel function or expression during pregnancy [24,36,37,40,41,43–74]. However, such data were sparse, so it was necessary to reference multiple data types across species with different gestation lengths. Unsurprisingly, we found that most available data were from a murine model with a 21-day gestation, but data from humans and guinea pigs were also available. As such, the timelines used in our study are all based on a 21-day gestation. Data types primarily consisted of mRNA data, although protein expression data and current density measurements were also used where available. Furthermore, this data was often contradictory, even within the same data type and species, with many datasets reporting different timing or causes of peak activity. For example, Inoue et al. described the change in total sodium current (I_Na_) during gestation as an increase in the fraction of cells expressing the channel [36]. Conversely, Yoshino et al. attributed it to an increase in current density across all cells [68]. With these considerations in mind, constructing a specific timeline was not possible, and we instead identified only the general magnitude of changes in ion channel activity and expression throughout pregnancy. Where data was lacking at multiple gestational time points, we compared changes from the non-pregnant to the pregnant state. Such was the case for the voltage-dependent potassium current (I_K1_). No gestational data was available for either the plasma membrane Ca^2+^-ATPase (PMCA) or the hyperpolarization-activated current (I_h_), and they were therefore assumed to remain constant. Data on current-voltage relationships over gestation, which are required to identify changes in gating properties, were available only for the L-type calcium current (I_CaL_), I_Na_, and the voltage-dependent potassium currents (I_K1_ and I_K2_) and showed no changes [36,64]. As a result, we assumed that the gating parameters for all currents, including the steady state values and time constants, also remained fixed throughout gestation. Other cell properties, such as the specific membrane capacitance (*C_m_* in Eq. (1)), were not reported to change significantly in the literature during late pregnancy and were therefore not modified [68].

#### 4.1.2. Sobol’ sensitivity analysis

Based on our literature review, we reduced gestational changes in ion currents to changes in their expression levels, which we modeled by scaling their maximal conductances (i.e., *g* in Eq. (2)). However, this still required fitting 14 parameters, and further reduction was necessary to ensure computational feasibility and facilitate convergence in the fitting process. As such, we performed a variance-based global sensitivity analysis of the Tong model to examine the effect of each current on relevant gestational EP characteristics, namely RMP and burst properties.

Currents were scaled by varying their maximal conductances between 0 and 2 times their baseline values. This upper value was chosen with reference to the general maximum gestational change in ion current conductances from our literature review. For the I_Na_ current, where a single baseline value is not defined, we scaled between 0 and 2 times the maximum range value reported by Tong et al. (0.125 nS/pF) [14]. The sodium-calcium exchanger current (I_NaCa_) was not tested because it depends on the J_NaCa_ flux and therefore lacks a maximal conductance parameter. At each iteration, the model was prepaced for 1,800 seconds without stimulus to steady state, at which point, the RMP was measured, and a 5-second stimulus of -0.4 pA/pF was applied. The properties of the resulting burst were automatically calculated using a variation of the method described in Reinl et al. [24] (see below).

The sensitivity of the Tong 2011 cellular model’s EP properties to each current was determined using a variance-based, global sensitivity analysis, introduced by Sobol’ [75]. Specifically, we used the implementation provided by the open-source Python library SALib to construct samples and compute first-order and total-order Sobol’ indices along with 95% confidence intervals [76,77].

Briefly, we begin by assuming that each measured property of the Tong model (i.e., the burst properties and the RMP) is described by a function of the form *Y* = *f*(*X*), where *Y* is the scalar output, and *X* is the vector of input parameters *X_i_*. The outputs are then normalized using their mean (μ) and standard deviation (σ) as *Z_i_* = (*Y_i_* ― μ)/σ. Assuming that all input parameters are independent and random, the first-order sensitivity index (*S_i_*) for the i^th^ input parameter is given by:

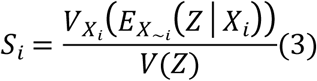

where *V* is the variance operator, *E* is the expected value operator, and the subscripts *X_i_* and *X*_∼*i*_ indicate that the operator is taken for only the parameter *X_i_* or for all parameters excluding *X_i_*, respectively. The first-order sensitivity index ranges from 0 to 1 and represents the normalized sensitivity of the output to the parameter *X_i_*. The total-order index (*S_Ti_*), which also ranges from 0 to 1, includes both the first-order effects of *X_i_* and its interactions with all other parameters, and is found using:

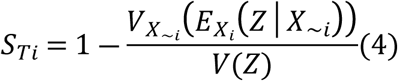

To estimate the sensitivity indices, we employed Saltelli’s method [78] to generate quasi-random sample points uniformly distributed in the parameter space. The Tong 2011 model was thus evaluated at each sample point, yielding estimates of the first- and total-order indices. For *N* = 4096 samples and *D* = 12 parameters, this resulted in *N*(*D* + 2) or 49,152 model evaluations. Confidence intervals were calculated using bootstrapping with a resample count of 100. Full details of the numerical methods used to estimate the Sobol’ indices can be found in the original works [75,78].

### 4.2. Single-cell model fitting

Based on the gestational regulation data and sensitivity analysis (Sec. 2.1), we chose to fit the I_CaL_, I_Na_, and I_K1_ currents. Although the model was highly sensitive to the I_NSCC_ current, literature showed that this current is not significantly regulated in late pregnancy [24]. Similarly, the I_Na_ current was chosen over the T-type calcium current (I_CaT_) as it showed more complex and significant regulation during late pregnancy [36,40,55,68]. Regarding the background current I_b_, although it had a substantial impact on many of the measured properties (Figure 1), it lacks a specific biophysical basis; instead, it represents a collection of minor potassium currents that lack sufficient electrophysiological data for inclusion. We therefore excluded this current from subsequent fitting analysis.

Using our reduced parameter set, we fit the Tong 2011 model to changes in RMP and burst properties characteristic of myometrial activation as given by Bengtsson et al. and Reinl et al., respectively [24,25]. Adopting the terminology used by Bengtsson et al., we chose three time points in late pregnancy of a rat model: days 16/17, 18/19, and 20/21 (term, before labor) [25]. For the Reinl dataset, which used a mouse model with a 19-day gestation, these roughly correspond to their day 14, 18, and 19 time points [24]. At each gestational time point, we employed MATLAB’s Bayesian optimization implementation (*bayesopt*) to fit the maximal conductances by minimizing the weighted mean-squared error (WMSE) of the RMP, the spike density, and the burst duration with weights of 0.5, 0.4, and 0.1, respectively. As the physiological stimuli in the uterus remain unknown, we also fit the stimulus amplitude (I_stim_) while assuming the stimulus duration equals the burst duration. Initial parameter ranges were set to 0.5 to 2 times baseline values for the maximal conductances and -0.1 to -0.5 pA/pF for the stimulus amplitude. However, these large ranges led to slow convergence and required manual tuning at each gestational time point based on preliminary Bayesian optimization runs. The final bounds on the maximal conductances at each gestational time point are given in Table 2.

**Table 2.** Bounds on maximal conductances used for Bayesian parameter estimation.

| Gestational Time Point | $g_{CaL}$ (nS/pF) | $g_{K1}$ (nS/pF) | $g_{Na}$ (nS/pF) | $I_{stim}$ (pA/pF) |
| --- | --- | --- | --- | --- |
| 16/17 | 0.3 – 0.6 | 0.572 – 0.832 | 0.0813 – 0.1375 | -0.1 – -0.2 |
| 18/19 | 0.45 – 0.72 | 0.52 – 0.676 | 0.0938 – 0.15 | -0.1 – -0.2 |
| 20/21 | 0.6 – 0.63 | 0.52 – 0.546 | 0.0938 – 0.15 | -0.1 – -0.2 |

At each iteration of the Bayesian optimization, the sampled maximal conductances and stimulus amplitude were set, and the resulting RMP and burst properties were measured. To account for the effects of the experimental conditions used in the literature sources, we set the model’s extracellular concentrations to those of the corresponding bath solutions (Table 3). As such, to measure the RMP, we first set the extracellular ion concentrations to those of the Bengtsson et al. bath solution [25], then measured the RMP as before. However, this caused random depolarizations and bursts when running the model for long durations, so we set I_Na_ to 0 for the RMP tests. Based on our sensitivity analysis (Figure 1), this should have had little impact on the final RMP value. Similarly, for the burst properties, we set the extracellular ion concentrations to the data from Reinl et al. [24], prepaced the model for 3 seconds, and measured the burst as before. The extracellular concentrations for each test are summarized in Table 3. Each optimization was run for an initial 90 iterations and continued in increments of 60 until it was deemed converged. Convergence was defined as a WMSE below 0.005 and tens of iterations without improvement.

**Table 3.** Extracellular concentrations of the original Tong model and bath solutions used in literature sources. The concentration values for our implementation of the Tong 2011 model were set to those of Bengtsson et al. [25] to measure the resting membrane potential (RMP) during Bayesian calibration, while values from Reinl et al. [24] were used to measure the burst properties. The original Tong model’s parameters were used for all other simulations outside of the Bayesian optimization.

| Source/Test | $[Ca^{2+}]_o$<br>(mM) | $[Na^+]_o$<br>(mM) | $[K^+]_o$<br>(mM) | $[Cl^-]_o$<br>(mM) |
| --- | --- | --- | --- | --- |
| **Original Tong** | 2.5 | 130 | 6 | 130 |
| **Bengtsson et al. (RMP)** | 2.5 | 143 | 5.9 | 128.7 |
| **Reinl et al. (Burst)** | 1.2 | 133 | 5.9 | 140.1 |

### 4.3. Excitation-propagation model

To extend our single-cell model to the tissue level, we employed a monodomain model to simulate the propagation of excitation (APs) between cells, which corresponds to the addition of an anisotropic diffusion term to Eq 1:

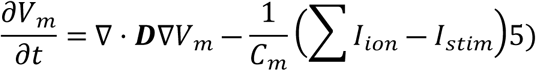

Assuming two fiber families, representing the longitudinal and circumferential muscle layers of the uterus [2], the anisotropic diffusion tensor ***D*** is defined as:

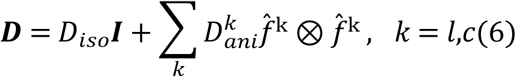

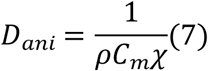

where ***I*** is the identity tensor, *D_iso_* is the isotropic diffusion coefficient, and 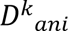 and *̂f^k^* are the anisotropic, fiber-oriented diffusion coefficient and unit direction vector of the k^th^ fiber family, respectively. The longitudinal diffusion coefficient 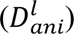 was found from Eq 7 using the specific resistivity value (*ρ*) for day 17-22 longitudinal rat myometrium from Sims et al. [30]. The surface area to volume ratio (*χ*) is calculated from Yoshino et al. [68]. As we did not find a specific resistivity value in the literature for the circumferential direction, the circumferential diffusion coefficient 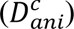 was derived by assuming 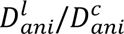 = 2.5. This ratio was chosen with reference to the longitudinal and circumferential conduction velocity ranges given in Rabotti & Mischi [7]. Finally, 5% of the diffusion tensor’s magnitude was heuristically set for *D_iso_*. Longitudinal fibers were assumed to lie uniformly in the y-direction and circumferential in the x-direction. As Sims et al. only provides a single resistivity value for late pregnant myometrium, the same diffusion tensor was used for all three gestational time points [30].

Single-spike and burst propagation properties were analyzed on a two-dimensional section of tissue measuring 100 mm x 100 mm. Single-spike APs were invoked using a stimulus of -5.0 pA/pF to a 5-10 mm square area in the bottom-left corner of the tissue, while bursting-type APs required a low-amplitude, long-duration stimulus (Fig. A2 of Supplemental Material). Tests to determine appropriate numerical parameters for the tissue model, such as the time step and mesh spacing, are described in Sec. S1 of Supplemental Material. Based on these results, we decided to use a mesh spacing range of 0.25 to 0.5 mm and a time step range of 0.5 to 1 ms for all subsequent tissue simulations.

### 4.4. Burst detection algorithm

We detected bursts in AP tracings using a method similar to that described by Reinl et al. [24]. Spikes with a minimum prominence of 7 mV were first identified using MATLAB’s *findpeaks* function and then grouped assuming a maximum interval of 1.5 seconds. Groups of spikes were classified as a burst only if their duration, defined as the time between the first and last spike, was at least 2 seconds; spike groups below this duration threshold were ignored. The maximum height and rate of rise of the first spike in the tracing were recorded, as well as the duration and spike density of the burst.

### 4.5. Model implementation

For the Sobol’ sensitivity analysis, we used the CellML [79] implementation of the Tong 2011 model. The model was solved in the OpenCOR environment [80], using the CVODE solver from the SUNDIALS library [81]. For the Bayesian optimization, the Tong model was reimplemented in MATLAB and solved using forward Euler.

Tissue-level AP conduction simulations were performed using an in-house multi-physics finite element solver, adapted from the open-source solver *svFSI* [82]. To evaluate the full reaction-diffusion equation given in Eq 5, we employed an operator-splitting method from Qu & Garfinkel [83] to decouple the integration of the ion currents from that of the diffusion term. In this method, Eq. (5) is rewritten as

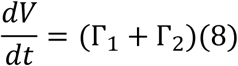

where Γ_1_and Γ_2_ represent differential operators, corresponding to the non-diffusion and diffusion parts, respectively. The integration of Eq 8 for one time step can then be approximated as

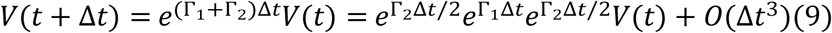

Therefore, at every time step, updating Eq. (5) proceeds as integration of the diffusion part for Δ *t*/2, integration of the ion currents for Δ*t*, and a final integration of the diffusion part for Δ*t*/2. The diffusion equation is integrated using the second-order implicit generalized-α method [84]. Gating variables were analytically integrated using the Rush-Larsen method [85], while the other state variables of the Tong 2011 model were integrated using the fourth-order Runge-Kutta method. All implementations were verified against the original Tong model [14] and the CellML version.

## Acknowledgments

The authors would like to acknowledge partial financial support from the National Science Foundation’s CAREER award (2443726) and from Columbia University’s Fu Foundation School of Engineering and Applied Sciences (SEAS) in performing this work.

## Supplemental Material

### S1. Tissue model numerical refinement tests

To identify appropriate simulation parameters, we conducted mesh and time step refinement tests. For the mesh refinement, the time step was held at 0.5 ms, while the mesh spacing was varied from 0.2 to 3 mm in steps of 0.2 mm. Similarly, for time step refinement, the mesh spacing was held at 0.5 mm, and the time step was varied from 0.2 to 4 ms in 0.2 ms increments. In both cases, a 10 mm tall area along the bottom edge of a 100 mm two-dimensional square domain was stimulated to produce a single-spike AP that propagated as a plane wave along the vertical axis. By producing a planar wavefront, we sought to isolate the effects of numerical parameters on conduction velocity from those of wavefront geometry and curvature, which have been shown to substantially affect AP propagation. The conduction velocity along the vertical axis was recorded and compared across time steps and mesh spacings (Fig. A1). Propagation failure was defined as a spike that failed to continue for more than 150 ms. To test the circumferential axis, the fiber directions were rotated 90 degrees, and the procedure was repeated.

The model showed high sensitivity to mesh spacing, especially in the circumferential direction, where conduction velocity rapidly declined from its peak at Δx = 0.4 mm. Propagation failure was then reached at Δx = 0.8 mm for the preterm time points and Δx = 1.2 mm for the day 20/21 time point. In contrast, the time step had a relatively low impact on the conduction velocity, with the minimum timestep for propagation failure occurring in the day 18/19 tissue at Δt = 2 ms. For the day 20/21 tissue, propagation failure never occurred within the measured time step range. Based on these results, we selected a mesh spacing range of 0.25 to 0.5 mm and a time step range of 0.5 to 1 ms for all subsequent tissue simulations.

### S2. Tissue stimulation protocol

**Figure A1.**
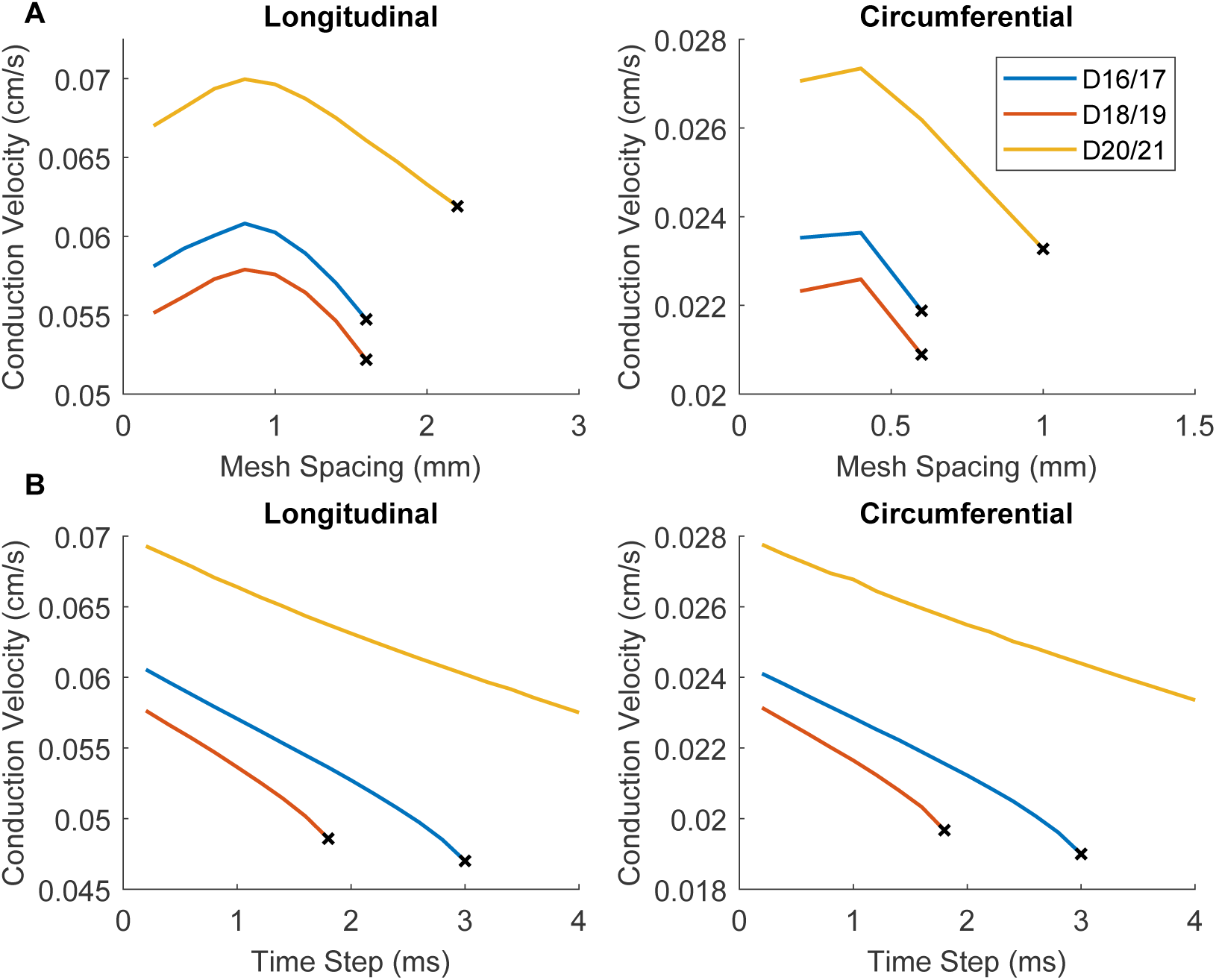
Mesh and time step refinement results. Effect of mesh spacing and time step on conduction velocity of a single spike along the longitudinal and circumferential directions. X marks the final data point before propagation failure. (A) Mesh refinement results were evaluated with the time step held at 0.5 ms. (B) Time step refinement results were evaluated with the mesh spacing held at 0.5 mm.

**Figure A2.**
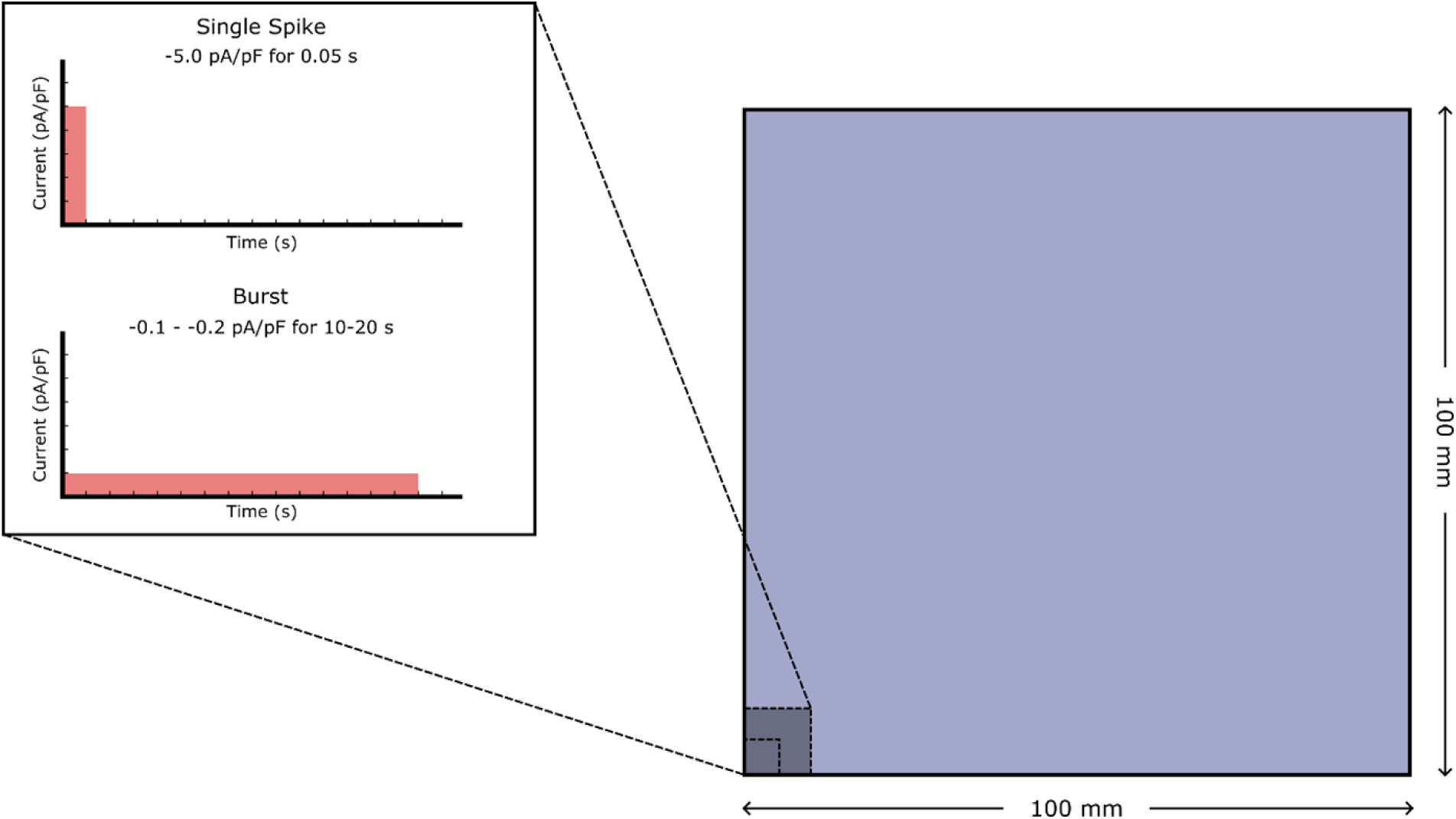
Tissue stimulation protocol. All tissue simulations were performed on a 2D domain spanning 100 mm x 100 mm. A stimulus current was applied to a 5-10 mm square region in the bottom left corner. The resulting AP morphology depended on the duration and amplitude of the applied stimulus. Single spike APs were elicited using a stimulus of -5.0 pA/pF for 50 ms, while a stimulus of - 0.1 to -0.2 pA/pF for 10,000 to 20,000 ms was used for bursting-type APs.

## Notes

### Competing Interest Statement

The authors have declared no competing interest.

